# Testing the mediating role of social interactions on the relationship between age and plumage traits on reproductive success

**DOI:** 10.1101/2025.03.06.641912

**Authors:** Zachary M. Laubach, Kayleigh P. Keller, Rebecca J. Safran, Toshi Tsunekage, Iris I. Levin

**Affiliations:** Department of Ecology and Evolutionary Biology, University of Colorado, Boulder, CO, USA; Department of Statistics, Colorado State University, Fort Collins, CO, USA; Department of Biology, Kenyon College, Gambier, OH, USA

**Author notes:** Correspondence: Zachary Laubach.

**Keywords:** age, barn swallow, causal inference, mediation, reproductive success, social network

## Abstract

Differential reproduction is a key driver of evolution that is determined by individual characteristics and mating opportunities, including mate choice. Social interactions between conspecifics are hypothesized to be important in facilitating mate choice and reproductive success but are difficult to measure. Using data from 52 adult barn swallows (*Hirundo rustica erythrogaster*), whose social interactions were measured via proximity tags, we tested the hypothesis that social interactions mediate the relationship between age (a proxy for experience) and condition-dependent plumage traits, and their associations with reproductive success. We found that older female barn swallows had higher fecundity and that older males have higher paternity. Older males achieved higher paternity through extra pair copulations, not by greater paternity with their social mate. Longer tail streamers were associated with greater fecundity/paternity in both sexes, but this effect was independent of age only among females. Darker ventral plumage coloration was not associated with higher reproductive success in either sex. We also observed that older birds appear to be less social with conspecifics, as indicated by fewer numbers of social interactions, though these associations were only marginally significant in males. Interestingly, females with fewer social interactions had higher fecundity. Finally, we found no evidence of mediation by the number of social interactions. Taken together, our results suggest that older, more experienced birds can produce more offspring while being less social.

## INTRODUCTION

Annual reproduction is a key component of Darwinian fitness that is determined by characteristics inherent to the individual and mating opportunities that are influenced by interactions with conspecifics [1]. Individuals that are in better condition or more experienced typically produce more and healthier offspring than animals in poor condition or lacking experience [2–5]. Condition and experience change with age and are often associated with expression of secondary sexual traits [6], and as a result, age and condition-dependent traits influence reproductive success. Reproduction is also influenced by social interactions, including intrasexual competition and intersexual mate choice [1]. Non-random mate choice based on traits that reliably convey information about partner attributes, including experience, is predicted to lead to higher reproductive success [7–9].

A mechanism through which individuals can sample and select a mate is social interactions. There are numerous examples wherein members of the opposite sex assess potential mates as part of their mate selection process. A conspicuous example is lek mating [10–12]. If social interactions allow individuals to assess potential mates’ traits, then more extensive interactions should optimize mate selection and increases reproductive success among more desirable mates [13].

We used data from North American barn swallows (*Hirundo rustica erythrogaster*) to ask: are age and/or plumage traits associated with reproductive success, and to what extent are these relationships mediated by female-male as well as same sex social interactions? Barn swallows are migratory passerines that breed in colonies of ∼1-50 pairs during the summer in North America. Social pairs typically produce 1-3 broods per breeding season, and each brood contains 3-5 eggs per brood [14]. In addition to mating within social pairs, approximately 41% of nestlings are sired through extra pair fertilization [15]. Reproductive success in this subspecies has been found to be positively associated with age [16–18] and experimental darkened ventral plumage color [19], as well as both positive [17] and negative [20] associations with tail streamer length.

Based on prior work in barn swallows, we hypothesized that older swallows as well as swallows with darker ventral plumage coloration would have higher reproductive success, and we were agnostic in our predictions about tail streamer length. We also hypothesized that older swallows would have darker plumage and longer tail streamers. Second, we hypothesized that older swallows and those with preferred plumage traits would have a greater quantity of social interactions. Third, we hypothesized that more social individuals would produce more offspring. Fouth, we hypothesized that the effect of age and plumage traits on reproductive fitness would be mediated by social interactions with group members.

To test these hypotheses, we conducted a mediation analysis. Briefly, mediation models involve decomposition of the total effect of the explanatory variable on the outcome into its direct and indirect effects [21,22]. The direct effect is the effect of the explanatory variable on the outcome that is independent of the mediator, while the indirect effect is the effect that passes through the mediator. The sum of the direct and indirect effects equals the total effect. To explore mediation, the total effect of age and/or plumage traits on reproductive success can be decomposed into the direct and indirect effects in scenarios involving varying strengths of mediation by social interactions (Figure 1, a-d). In the context of our hypotheses, complete mediation would involve a scenario in which the effect of age or plumage traits on reproductive success is entirely dependent on social interactions, such that the parameter estimate for age or plumage is attenuated to the null when conditioning on the social interaction.

**Figure 1.**
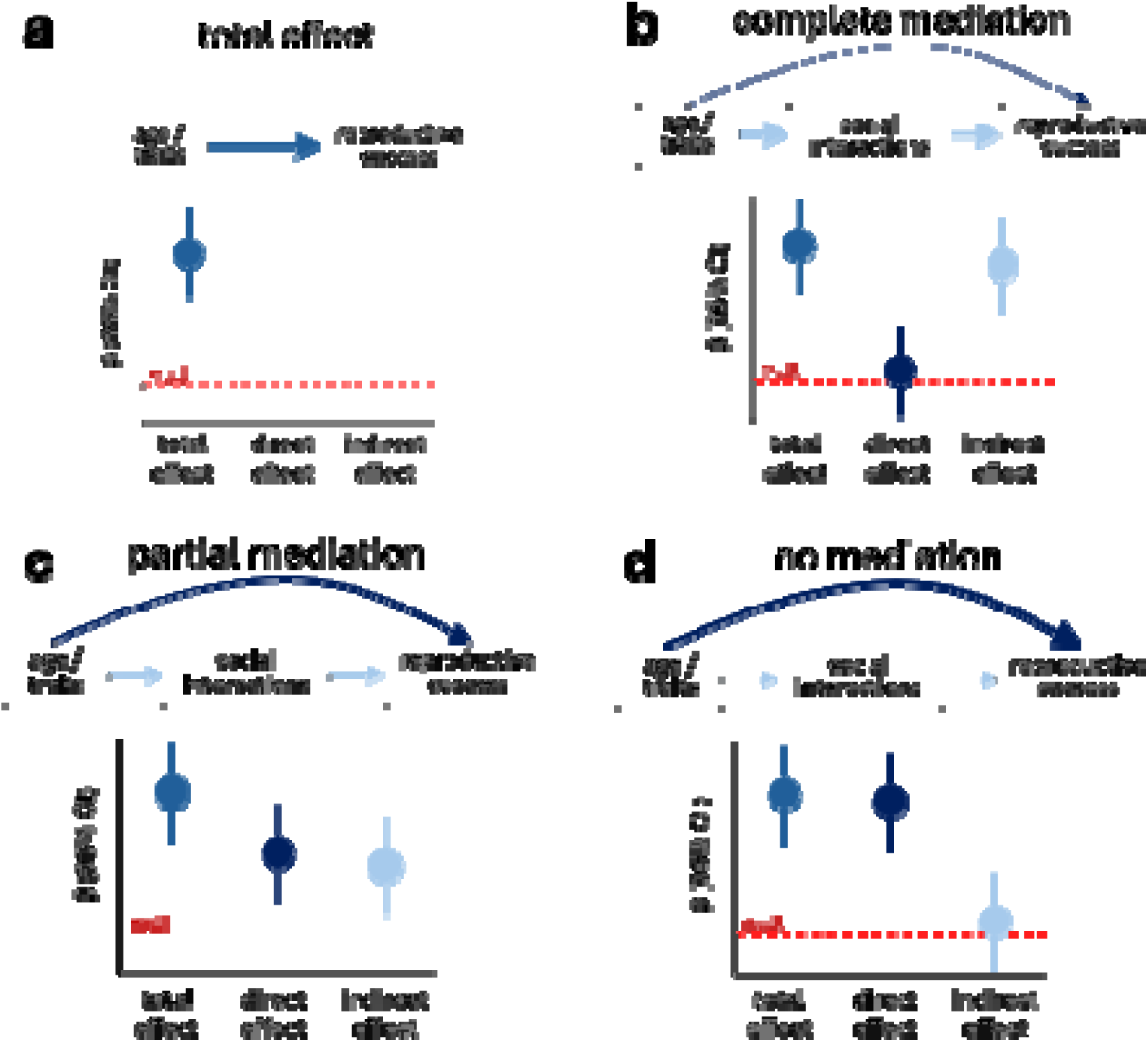
Directed Acyclic Graphs (DAG) of the hypothesized biological mechanism in which age and/or morphological traits affect reproductive success, and social interactions mediate these relationships. The predicted decomposition of **a)** the total effect into the direct and indirect effects under scenarios of **b)** complete, **c)** partial, or **d)** no mediation. Total effect = direct effect + indirect effect.

## METHODS

### Study population

We studied 61 wild adult barn swallows in a large building at a horse rescue facility during the breeding season by monitoring nests between 1 May – 9 September 2015 in Boulder County, CO, USA. Using mist nets, we first captured 58 of the 61 (95%) adult birds on June 11 or June 13 and fit each bird with a U.S. Geological Survey numbered aluminum leg band and a unique combination of 2 plastic color bands. We determined the sex of each bird as part of the capture procedure. Female birds were identified based on the presence of a brood patch, and generally exhibiting lighter ventral plumage and having shorter wing and tail streamer lengths compared to males. The sex of each member of a social pair was confirmed through behavioral observations at their nest. We determined a bird’s age based on previous banding records.

Given the near complete banding at the site in previous years and the strong breeding philopatry, any unbanded birds were assumed to be first time breeders. We dichotomized continuous age as a two-level categorical measure corresponding to ‘second year’ (SY) birds in their first breeding season, and ‘after second year’ (ASY) which included any birds with prior breeding season experience.

During the initial capture we also measured morphology, collected feathers for plumage assessment, sampled blood for paternity analyses, and attached a proximity logger for quantifying social interactions (Figure 2). Shortly after the initial capture and before collection of any social interaction data, on June 12, we collected the first clutch of eggs from 26 active nests. This was necessary to synchronize breeding phenology so that we could compare social interactions among all adults in the colony. Based on the clutch initiation dates, we determined that 24 of these nests included complete clutches, while for 2 nests, both containing 3 eggs at the time of collection, we could not determine if females had completed egg laying. On the morning of June 13, all proximity loggers were turned on and remained active through June 15, which we refer to as the pre-manipulation period. Next, a random sample of males were captured the night of June 16 off or near their nests to have their ventral feathers experimentally darkened. Proximity loggers were reactivated on June 19 and continued collecting data until they were removed from the bird or the battery died. Females laid eggs for up to three total clutches including the first clutch that we collected. Similarly, males contributed paternity in up to three clutches, however due to incomplete paternity from eggs (not all embryos were developed enough for DNA extraction) we restricted male reproductive success to clutches 2 and 3. Second clutches were initiated between June 20 – July 15, and third clutches were initiated on or after July 26. Second and third clutch nestlings had blood drawn from their brachial vein (∼ 30uL in capillary tubes) 12 days after hatching, and we collected blood samples from adult birds as part of our capture protocol and to determine relatedness.

**Figure 2.**
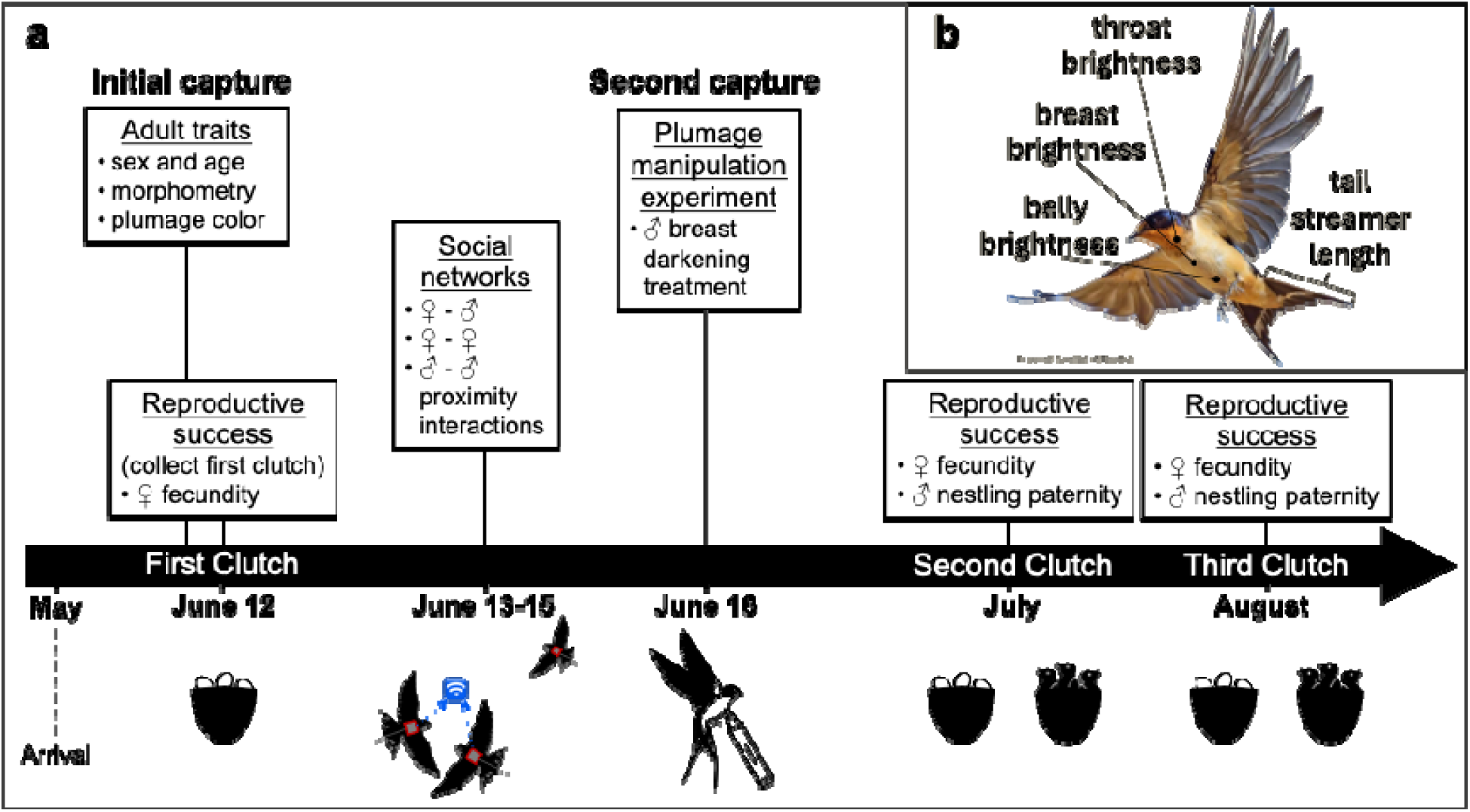
**a)** Overview of the complete data collection timeline and experimental manipulation. **b)** This inset shows the plumage traits that we measured which have previously been found to be associated with reproductive success in this species. Swallow vector images is from the Noun Project and swallow photograph from Cornell Lab – All About Birds.

### Trait assessment and experimental manipulation

To measure plumage color, we plucked 15-20 feathers sampled from three regions of the ventral surface (throat = T, breast = R, belly = B) that were previously shown to be associated with reproductive success and/or mate assortativity [19,23] (Figure 2, b). Feathers were stored in opaque envelopes until feather brightness was quantified via spectrometry. We used an Ocean Optics USB4000 spectrometer (Dunedin, FL) and a pulsed xenon light (PX-2, Ocean Optics) to measure throat, breast, and belly brightness. Brightness is a measure of light reflectance from a surface, and as such that darker plumage has a lower spectral signal. We arranged feathers in an overlapping fashion on a white index card, like their natural orientation on a live bird [23]. Then a 300-700nm spectrometry probe, positioned perpendicularly and at a set distance, was used to sample each set of feathers with 20 scans across the feathers and during three separate sampling iterations. We focused on reflectance values corresponding to the red/brown region of the light spectrum (600-700nm). Spectral measures for each body region were averaged and calibrated against a white standard (Ocean Optics WS-1) and a dark standard (all light excluded) using SpectraSuite v.3.0.151 (Ocean Optics). In addition to measuring both female and male natural plumage brightness, we also quantified male breast brightness after manipulating ventral plumage color using a PrismaColor Light Walnut (#3507) marker to artificially darken the ventral feathers of 14 randomly assigned males; see [24].

### Social interactions

Social interactions between barn swallows were recorded from Encounternet proximity loggers, which are small tags, approximately 1.2 g each, that send and receive unique radio signals at 433MHz [25,26]. Proximity loggers record non-directional association counts as well as durations of time during which tags are within close proximity to each other. Previous work has shown that social interactions measured from these tags are related to plumage traits, physiological changes and microbe transmission [24,26]. To quantify social interactions using the proximity loggers, we partitioned raw data into three subsets, one including female-male interactions and two for same sex interactions (*i.e.*, female-female and male-male). These interactions were further filtered to include a date range during which all proximity loggers were simultaneously active, and to include encounters at or less than a maximum distance of ∼0.1m based on filtering the minimum average receiver signal strength indicator (RSSI) values to ≥ 20 [25]. These data include a pairwise count of each proximity encounter and the duration of time that birds remained in close proximity. Tags were set to collect interaction data from 29 female and 29 male barn swallows between approximately 6:00 - 9:00 am before and after the plumage manipulation experiment. Pre-manipulation interaction data were collected between June 13-15, while post-manipulation data were available from June 19-20. Pre- and post-manipulation adjacency matrices were used to calculate node level social network metrics. We quantified degree, the total number of unique group mates that an individual interacts with, and strength, the sum of the total number of interactions [27].

Several of the originally tagged birds (n=58) were excluded from our final pre-manipulation and post-manipulation analyses. One female left the site and two additional females lacking complete social interaction data were excluded from analyses (n = 26 adult females). Similarly, two males were excluded because they lacked reproductive success data, and another male died or left the site (n = 26 adult males). When calculating degree and strength, we retained the social interaction data for birds with incomplete reproductive success data before removing them from the final data sets because they contribute to the strength and degree of other birds. Therefore, we had social interaction data on 88% of the resident adults at our study site.

### Reproductive success

We monitored nests every 3 days throughout the breeding season to measure reproductive success. We counted the total number of eggs each female laid (fecundity) for up to three clutches over the duration of the breeding season. For males, we measured reproductive success by estimating his total paternity and dividing total paternity into each males’ within pair paternity (WPP; offspring sired with his social mate) and extra-pair paternity (EPP). Paternity was determined using DNA from ∼ 12 day old nestlings. Blood was stored in lysis buffer until DNA was extracted using a DNeasy Blood and Tissue Kit (Qiagen, MD, USA, Cat No./ID: 69506). Six microsatellite loci (Escu6, Hir11, Hir19, Hir20, Ltr6, and Poc6) were used to assign paternity of offspring following Jenkins et al. 2014 [28]. We genotyped adults and nestlings using PeakScanner (v2.0, Applied Biosystems, CA, USA) and used CERVUS (v3.0, [29]) to assign paternity. Our combined first-parent exclusion probability was 0.993.

We quantified female reproductive success based her fecundity from all three nesting attempts. First nesting attempts were not synchronized, and as result we were only able to extract DNA from 21 of 27 active nests. Due to incomplete sampling, assessment of male reproductive success was limited to second and third broods. For the second and third nest attempts, we extracted DNA from nestlings that survived until day 12 from all active nests and assigned paternity to 91% of those nestlings.

### Statistical analysis

To test our hypotheses, we implemented a causal mediation analysis that extends the conditional linear regression approach through the estimation of natural direct and indirect effects, even in the presence of an interaction between the explanatory and mediator variables [30–32]. This entailed four primary steps, 1) establishing a relationship between explanatory and outcome variables, 2) estimating the association between the explanatory and mediator variable, 3) quantifying the effect of the mediator on the outcome variable, and 4) fitting a model that includes the explanatory, mediator, and outcome variables using the causal mediation functions in the ‘*mediation*’ package to estimate the natural direct and indirect effects [33].

First, we used linear regression to estimate the associations between each pair of explanatory and outcome variables, as well as the association between age and plumage traits for females and males separately. Among females, we began by regressing total fecundity on age, a two-level categorical variable. We ran separate models in which standardized values of plumage traits, throat, breast and belly brightness as well as right tail streamer length, were explanatory variables and total fecundity was the outcome. For males we ran similar models as for females, with the notable distinction that here we assessed the association between combinations of male age and each plumage trait with total paternity, EPP and WPP as outcomes in separate models. We also modeled the effect of ventral plumage brightness after the experimental darkening treatment. For both sexes, we assessed the association between age and plumage brightness. Finally, where any plumage trait was associated with reproductive success, we also assessed the effect of plumage independent of age by including age as a covariate in those models.

Second, we sought to establish whether there was any association between the explanatory variable and potential mediator variable. We used linear regression to estimate the associations between age and plumage traits as explanatory variables with social interaction values, quantified as strength and degree, as outcome variables. Only plumage traits that were associated with reproductive success were carried forward into these models and we adjusted for age as confounder of the relationship between plumage traits and social interactions. Strength and degree were quantified from female-male, female-female, and male-male interaction networks.

Third, we ran linear regression models to assess the associations between the social interactions and reproductive success to understand the relationship between mediator and outcome. Finally, we proceeded with the causal mediation models, which we implemented using the mediate() function in the ‘*mediation*’ package in R [33]. Here we restricted our analyses to include variables that met the criteria for mediation, namely that the explanatory variable was associated with the outcome and mediator and the mediator is associated with the outcome. This included age and plumage traits that were associated with reproductive success, the outcome of interest. The subset of mediation models was further restricted based on associations between age and plumage traits with social interactions as well as social interactions with reproductive success. We tested for interactions between our explanatory variables and mediators in models where reproductive success was the outcome and included interactions terms in our final mediation models if the estimates for the interaction terms were statistically significant.

All data management and analyses were conducted in in R, version 4.4.2 (R Core Team 2024) using ‘*tidyverse’* [35], and ‘*igraph’* packages [36,37]. In all models, α = 0.05 was set as the threshold for statistical significance. Given our modest sample size, we balanced our inference and decisions about which variables to carry forward in subsequent analysis steps based on the consistency, direction, and magnitude of effect estimates alongside of conventional significance thresholds. We focused our reporting and interpretation of results based on estimates of effect size and confidence intervals, where confidence intervals that overlap the null indicate a lack of statistical significance at α = 0.05. However, due to moderate samples sizes, we also cautiously discuss estimates for a relaxed statistical significance threshold based on an α = 0.10.

### Ethical note

Research protocols involving the use of wild barn swallows were approved by the University of Colorado’s IACUC (permit no. 1303.02), and all experiments were performed in accordance with relevant guidelines and regulations. In addition, this work was conducted under the Bird Banding Lab, master permit bander ID 23505. Capture using mist nests in Boulder, Weld and Jefferson Counties, Colorado, was approved by the Colorado Parks and Wildlife issued under permit license number 15TRb2005. To generate complete social interaction networks and paternity data, we focused on capturing and tagging all adult birds at a single large breeding site as described above. To minimize stress, birds captured in mist nets were immediately removed, transported in opaque cloth bags to a nearby safe location, processed (see prior sections of the methods), and immediately released. We collected eggs from first clutches as early as possible to generate complete measures of fecundity and paternity while allowing plenty of time for birds to complete multiple additional clutches during the breeding season.

Egg collection was also necessary to synchronize the reproductive cycles of birds to accurately measure social interactions, thus avoiding a situation in which some birds were searching for mating opportunities while others were incubating or caring for nestlings. The backpack style proximity loggers balanced the costs of weight (1.2 g) with the need to house of a sufficient power source for data collection at two specified time periods over several days of the breeding season. Birds were captured after the experiment and proximity loggers were removed. Finally, for the plumage manipulation experiment, we used a non-toxic permanent marker.

## RESULTS

### Background characteristics

Among 26 female and 26 male barn swallows included in our final data, 23% were second year (SY) females, 27% after second year (ASY) females, 23% SY males, and 27% ASY males (Table S1). On average, older females had darker ventral plumage and longer tail streamers than young females, older males had darker ventral plumage and longer tail streamers than young males, and males had darker ventral plumage and longer tail streamers than females (Table S2). In female-male social networks, average SY female strength was 51.08 (± 21.19) and ASY female strength was 39.21 (± 18.31) interactions, and average degree was 14.33 (± 6.08) and 12.07 (± 5.11) unique social partners for SY and ASY females, respectively. Also, from female-male social networks, SY males had an average strength of 49.08 (± 26.98) and ASY male strength was 34.29 (± 15.11), and average SY male degree was 12.67 (± 5.43) and ASY male degree was 11.79 (± 4.61). In same sex social networks average SY female strength was 42.75 (± 23.57) and ASY female strength was 33.43 (± 23.57) social interaction, while SY male strength was 29.83 (± 18.75) and ASY male strength was 26.79 (± 13.55) (Table S3). For same sex social network measures of degree, younger females had on average 3 more unique female social partners than older females, while older males had on average 1 additional male social partner than younger males (Table S3). Average total fecundity was 8.50 (± 2.91) and 11.00 (± 3.04) eggs among SY and ASY females, respectively (Table S4). The average number of total offspring sired by SY males was 2.92 (± 2.11) and by ASY males was 6.00 (± 3.88). Similarly older males had higher average EPP (4.29 [± 3.24]) than younger males (1.50 [± 1.38]) while average WPP for ASY males was 1.71 (± 1.38) and for SY males was 1.42 (± 1.38).

### Step 1: Trait associations with each other and reproductive success

Older females produced 2.50 (95% CI: 0.08, 4.92) more eggs than younger females (Table 1). Similarly, older males consistently sired more total offspring and had higher EPP than younger males (total offspring, 3.08 [95% CI: 0.49, 5.68] and EPP, 2.79 [95% CI: 0.70, 4.87]. There was no difference in WPP by male age (Table 1). and over the duration of all three nesting attempts 5.91 (95% CI: 2.79, 9.02 eggs/nestlings). Females with longer tail streamers had higher fecundity even after adjustment for age, 1.59 (95% CI: 0.25, 2.93), but none of the female ventral plumage traits were associated fecundity (Table 1). Among males, their longer tail streamers were associated with higher total paternity and higher EPP, but these effects were attenuated to the null after adjustment for male age. Similarly, darker belly plumage was associated with higher WPP, but again adjustment for age attenuated this effect (Table 1). No other male plumage traits, including experimentally darkened ventral color were associated with any measure of male paternity (Table 1). Age was positively associated with female (4.79 [95% CI: 1.23, 8.35] mm) and male tail streamer length (4.36 [95% CI: -0.86, 9.58] mm), albeit the association in males was only significant at P<0.1. There was not association between age and ventral plumage color in either sex (Table S5).

**Table 1. Associations of barn swallow age and plumage traits with reproductive success.**
|  | female reproductive success<br>(N = 26) | male reproductive success<br>(N = 26) |  |  |
| --- | --- | --- | --- | --- |
|  | nests 1-3 fecundity | nests 2-3 total<br>paternity | nests 2-3 EPP | nests 2-3 WPP |
| | $\beta$ (95% CI) | $\beta$ (95% CI) | $\beta$ (95% CI) | $\beta$ (95% CI) |
| **categorical age** |  |  |  |  |
| second year | 0.00 (Reference) | 0.00 (Reference) | 0.00 (Reference) | 0.00 (Reference) |
| after second year | 2.50 (0.08, 4.92) | 3.08 (0.48, 5.68) | 2.79 (0.70, 4.87) | 0.30 (-0.82, 1.42) |
| **plumage traits \*** |  |  |  |  |
| throat brightness | -0.82 (-1.93, 0.70) | 0.28 (-1.19, 1.75) | 0.11 (-1.11, 1.32) | 0.17 (-0.40, 0.74) |
| breast brightness | -0.43 (-1.78, 0.90) | -0.94 (-2.38, 0.48) | -0.81 (-1.80, 0.57) † | -0.33 (-0.88, 0.23) † |
| belly brightness | -0.12 (-1.48, 1.22) | -0.60 (-2.05, 0.85) | -0.07 (-1.26, 1.15) | -0.53 (-1.68, -0.41) |
| belly brightness - age adjusted ‡ | - | - | - | -0.54 (-1.11, 0.03) |
| tail streamer length | 1.79 (0.65, 2.93) † | 1.37 (0.01, 2.73) | 1.24 (0.14, 2.34) | 0.13 (-0.44, 0.70) |
| tail streamer length - age adjusted ‡ | 1.59 (0.25, 2.93) † | 0.65 (-0.42, 2.32) | 0.88 (-0.22, 1.95) | - |

\* Plumage traits are z-score standardized and each trait is assessed in separate models.
† When plumage traits were associated with reproductive success, sex males from a second model that controlled for age are reported.
‡ Same as † but females in these models due to missing tail streamer length measurement to account for confounding by age.
§ We ran sensitivity analyses including the treatment effect, first as a continuous measure of a males breast plumage brightness after experimental darkening, and a separate model in which a categorical variable for received color manipulation, yes vs. no, was included. Estimates for breast plumage brightness were similar across all models.
Among males, total paternity includes nestlings from males sired with their social mate plus those sired via extra pair paternity, i.e., males total paternity, while EPP = extra pair paternity and WPP = within pair paternity. Male paternity from the first nest attempt is not included because paternity assignment from eggs collected was incomplete.
Beta sex males (95% CI) are from linear models.
Beta sex males are significant at $P < 0.05$ , and italicized estimates are significant at $P < 0.10$ .

### Step 2: Trait associations with social interactions

In both females and males, older birds appeared to have lower interaction strength in female-male networks, -11.87 interactions (95% CI: -27.05, 3.31) and -14.80 interactions (95% CI: - 31.29, 1.69), respectively, although this association was only significant in males at a relaxed P<0.1. Focusing on plumage traits that were associated fecundity, females with longer tail streamers had lower strength (-8.98 [95% CI: -18.15, 0.19] interactions per 1 SD tail length) and degree (-2.18 [95% CI: -4.73, 0.37] social partners per 1 SD tail length) from female-male networks independent of age, but again this association was only significant at P<0.1. Female and male age were not associated with strength or degree from same sex social networks. Focusing on male plumage traits that were associated with reproduction, male tail streamer length and belly brightness were not associated with strength or degree from either opposite or same sex networks (Table 2).

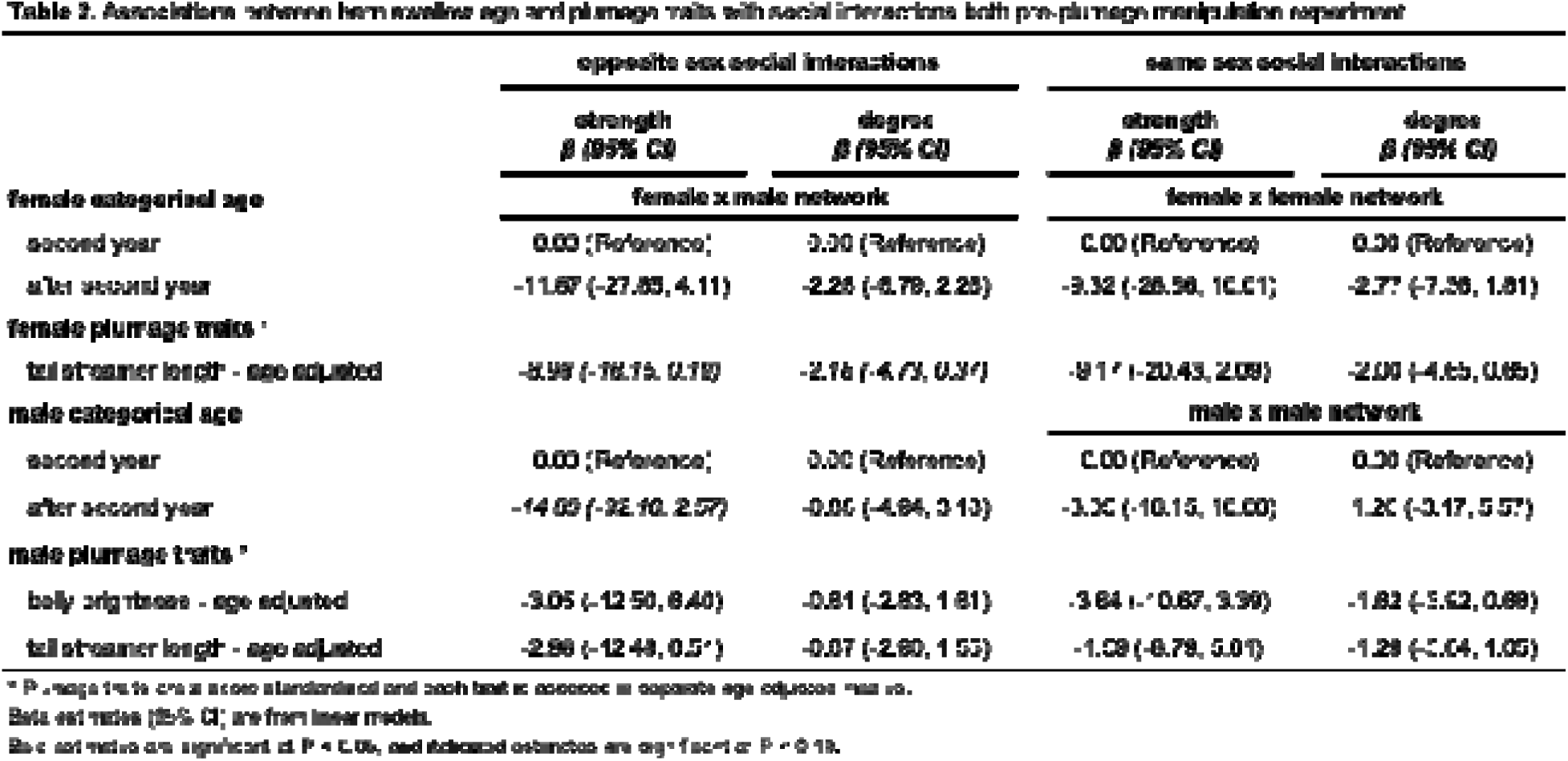

### Step 3: Social interaction associations with fitness and exploring mediation assumptions

Female barn swallows that more frequently interacted with males had lower fecundity (-1.33 [95% CI: -2.55, -0.11 eggs per 1 SD strength). In female-female interaction networks, females with higher strength also laid fewer eggs (-1.09 [95% CI: -2.35, 0.18]; eggs per 1 SD strength; Table 3), although this association was only significant at P<0.1. We observed no association between female degree and fecundity. We also observed no relationship between male interaction strength or degree from either opposite sex or same sex networks with any measures of reproductive success (Table 3).

**Table 3. Associations between barn swallow social interactions and reproductive success.**
| social interactions | female reproductive success | male reproductive success |  |  |
| --- | --- | --- | --- | --- |
|  | nests 1-3 fecundity | nests 2-3 total paternity | nests 2-3 EPP | nests 2-3 WPP |
| | $\beta$ (95% CI) | $\beta$ (95% CI) * | $\beta$ (95% CI) * | $\beta$ (95% CI) * |
| **female x male network** |  |  |  |  |
| strength | -1.33 (-2.55, -0.11) | -0.12 (-1.80, 1.35) | -0.41 (-1.81, 0.78) | 0.29 (-0.27, 0.85) |
| degree | -0.88 (-2.00, 0.62) | 0.75 (-0.88, 2.18) | 0.42 (-0.78, 1.62) | 0.33 (-0.23, 0.88) |
| **female x female network** |  |  |  |  |
| strength | -1.09 (-2.35, 0.18) | - | - | - |
| degree | -0.97 (-2.25, 0.31) | - | - | - |
| **male x male network** |  |  |  |  |
| strength | - | 0.44 (-1.02, 1.90) | 0.02 (-1.20, 1.23) | 0.42 (-0.12, 0.97) |
| degree | - | 0.54 (-0.91, 2.00) | 0.20 (-1.02, 1.41) | 0.35 (-0.21, 0.90) |

\*Because network variables are z-score standardized and each variable is assessed in separate models.
Among males, total paternity includes nestlings that males sired with their social mate plus those sired via extra pair paternity, i.e., males total extra-pairity, while EPP = extra pair paternity and WPP = within pair paternity. Male paternity from the first nest at arrival is not included because paternity assignment from eggs collected was incomplete.
Beta estimates (95% CI) are from linear models.
Beta estimates are significant at $P < 0.05$ , and italicized estimates are significant at $P < 0.10$ .

### Step 4: Causal mediation by social interactions

Based on evidence from steps 1-3 of our analysis, we ran causal mediation models focusing on female age and female tail streamer length as explanatory variables, female strength from female-male interaction networks as the potential mediator variable, and fecundity as the outcome. We found no significant interactions between both female age strength as well as female tail streamer length and strength in models where fecundity was the outcome Table S6), so we proceeded without further consideration of the potential mediator also acting as an effect modifier. In the causal mediation model involving female age, strength from female-male interaction networks, and total fecundity, the total effect of older age was 2.48 (95% CI: -0.14, 5.22) more eggs, the natural direct effect of older age was 1.86 (95% CI: -0.43, 4.26), and the natural indirect effect of age through strength was 0.61 (95% CI: -0.23, 1.88; Figure 3, a; Table S7) more eggs. In the mediation model involving female tail streamer length, strength from female-male interaction networks, and total fecundity, the total effect of longer tail streamers was 1.59 (95% CI: 0.17, 3.06) more eggs, the natural direct effect of longer tail streamers was 1.35 (95% CI: -0.31, 3.24), and the natural indirect effect of tail streamers through strength was 0.24 (95% CI: -0.44, 1.16; **Figure 3, b; Table S7**) more eggs. We did not run causal mediation models involving male age or plumage traits, any social network measures from opposite or same sex networks, and any measure of male reproductive success because assumptions for these models were not met in steps 1-3 as described above.

**Figure 3.**
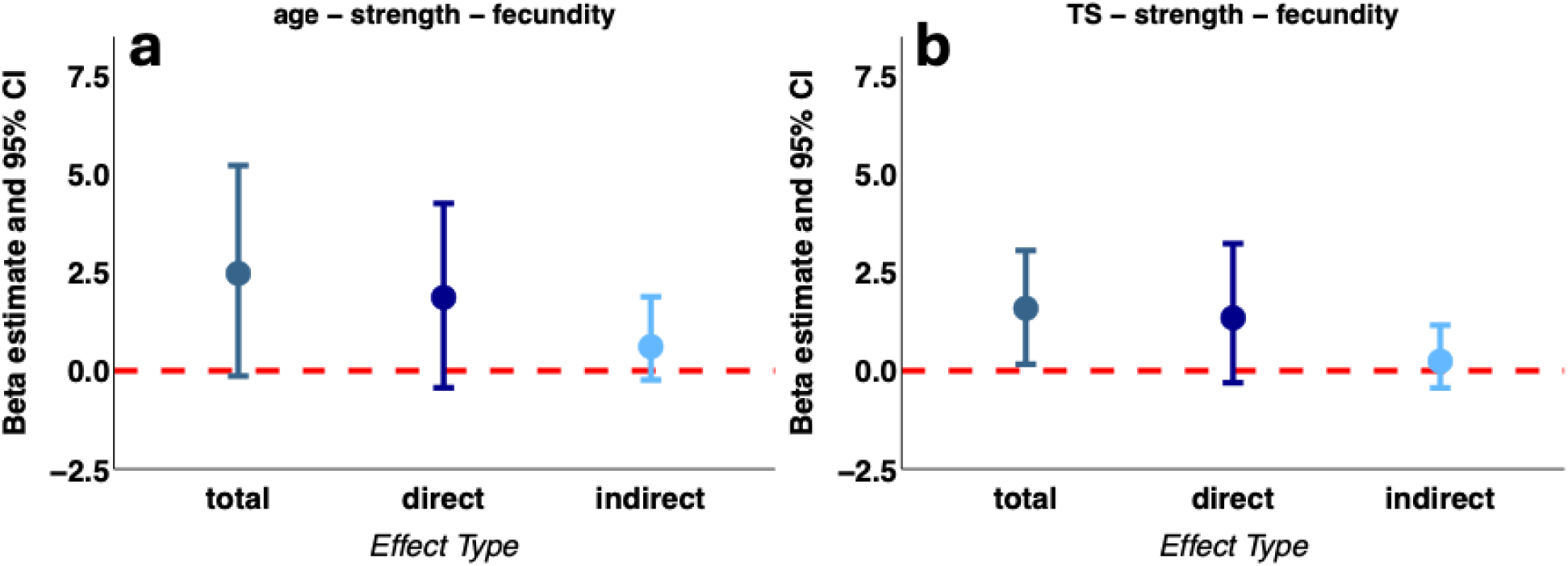
Estimates from the female barn swallow causal mediation models for **a)** the effect of age on fecundity and mediation by strength from female-male interaction networks and **b)** the effect of tail streamer length on fecundity and mediation by strength from female-male interaction networks. Estimates include the total effect estimated from each model and its decomposition into the natural direct effect (aka, average direct effect; ADE) and the natural indirect effect (aka, average causal mediation effect; ACME).

## DISCUSSION

We conducted a mediation analysis to determine if age and plumage traits affected reproductive fitness and the extent to which those effects were mediated by social interactions. Using data from 52 adult barn swallows that co-occupied a single breeding site, we found that older females produced more eggs. Females with longer tail streamers also had higher fecundity, independent of age. For males, older age was associated with greater total paternity, a pattern the was driven by older males achieving higher EPP but not WPP. In both sexes, darker ventral plumage was not associated with greater reproductive success, aside from darker male belly plumage being associated with greater within partner paternity. We found some weak evidence that older birds were less social, as indicated by lower strength from female-male networks, though these associations were only significant in males at a relaxed alpha threshold. Similarly, females with longer tail streamers had lower strength in opposite sex networks, independent of age, but again at a relaxed alpha. Interestingly, females but not males with lower strength from both opposite and same sex networks had higher fecundity, albeit with a confidence interval slightly overlapping the null when strength was from same sex networks. Finally, evidence of mediation by social interactions was lacking.

### Age and tail streamer length affected fitness

In our population, older birds with at least one or more prior years of breeding experience had higher annual reproductive success than first time breeders. Older females had higher fecundity, while older males had higher total paternity, a pattern that reflected their higher rates of EPP. An increase in annual reproductive success with increasing age prior to senescence is common among many passerine species [38–42], including in barn swallows [17,18,43,44].

Moreover, our finding of higher rates of EPP among older males is consistent with previous work in barn swallows [45]. Two non-mutually exclusive hypotheses that might explain these results, include 1) that individuals improve their reproductive competence or fertility with age [18] or 2) there is selective disappearance of certain phenotypes at the population level [46].

According to the competence hypothesis, more experienced individuals are better able to locate and secure resources, including nest sites, food, or mates. In an experimental study of captive zebra finches (*Taeniopygia guttata)* more experienced females laid more eggs and initiated their clutches earlier than inexperienced females [47]. Older male barn swallows are also hypothesized to increase their annual reproductive success by breeding earlier than younger males [18]. While the explanation that older birds are more experienced may extend to our results, whether higher reproductive success was due to experience *per se* or some other age-associated residual confounding remains unknown in our population. Nevertheless, greater experience leading to improved reproductive outcomes independent of age has been found in several wild birds, including both male and female Seychelles warbler (*Acrocephalus sechellensis*) [48], female collared flycatchers (*Ficedula albicollis*) [5], and western gulls (*Larus occidentalis*) [49].

In addition to experience, individuals that are in better condition typically have higher reproductive success [50,51], since they can invest more resources in reproduction [52–54]. Information about condition is often conveyed through expression of signals that are displayed during social interactions and used to assess potential mates [7]. In North American barn swallows, both tail streamer length and ventral plumage coloration are thought to be age/condition dependent signals [15,16]. There is also mixed evidence that these plumage traits are sexually selected signals since males with darker ventral plumage [19,23] have higher reproductive success, and males with both shorter [20] and longer tail streamers also having higher reproductive success [17]. Here, we found that longer tail streamers were associated with higher fecundity in females, and both higher total and extra pair paternity in males.

However, an effect of tail streamer length on reproduction independent of age was only evident in females and was confounded by age in males in our population as has previously been shown [18]. Given that both results from this study and prior work have found that tail streamer length increases with age in female [16] and male [17,18] barn swallows, trait covariation with age and reproductive success is expected. Notably, older male barn swallows appear to obtain higher rates of paternity both by enhanced fertility and breeding earlier [18]. Because we focus on paternity from second and third clutches after collecting eggs from first clutches, we minimize the advantage of earlier breeding among older males, which may further hint that older males are in better condition than young males.

We also assessed natural plumage coloration in both sexes, as well as experimentally darkened ventral plumage in males, and only found that males with naturally darker belly plumage had higher WPP but not total paternity. While this may suggest female mating preference for darker social mates, the role of darker ventral plumage as a sexually selected trait was not evident from our analyses. A lack of association between ventral coloration and male total paternity might partially be explained by the synchronized breeding in this study, and its effects on how birds prioritized within pair vs. extra pair mating. We also did not find that older birds had darker ventral plumage. Contrary to our results, other studies have found age related change in the expression of plumage color signals that influence reproductive success [55,56], and previous work in North American barn swallows indicates that darker ventral plumage coloration is under sexual selection [19,23]. Our null findings may reflect a context dependence that decouples the signal from age related condition and fitness [57–60], and that mate selection based on trait expression is dynamic across years and environmental conditions.

An alternative to the competence hypothesis, selection pressure imposed by migration may have removed poor-condition individuals from our population as has been found in European barn swallows [61], thus confounding the relationship between age and reproductive success. Still, we have previously found that females increased their reproductive success and increased expression of condition dependent plumage traits as they age [16]. Empirical support for both the improvement of individual competence [62] and the selection hypotheses [63] have been found in studies of wild birds but require longitudinal data to disentangle population from individual level effects [40,64]. If an age-related increase in individual performance was coupled with selective removal of poor-condition swallows from our study population, then our estimate for the effect age on reproductive success may be biased away from the null.

### Age and female tail streamer length were weakly associated with opposite sex social interactions

We found that older swallows tended to have lower strength when interacting members of the opposite sex. Older males had on average ∼9 fewer social interactions and older females had ∼12 fewer social interactions than younger birds, though 95% confidence intervals overlapped the null in both sexes. Similarly, we found a weak association in which females with longer tail streamer length had few social interactions with males, independent of age. Neither age nor plumage traits were associated with strength or degree from same sex social interaction networks.

Several recent studies have found a decrease in social interactions with age that was attributable, in part, to within individual change in social behavior and not just selective disappearance. In griffon vultures (*Gyps fulvus*) older individuals had lower interaction strength [65], a finding like in forked fungus beetles (*Bolitoherus cornutus*) that found experience decreased strength and betweenness with increasing age [66]. Older, more experienced, or birds in better condition might have been more selective in their social interactions, or they prospected less for mating opportunities. Still, demographic processes, including the appearance and disappearance of individuals and changes in population age structure, can influence the pool of individuals available for interactions [67–69], and contribute to age related differences in social connectedness, which we cannot rule out.

### Social interactions did not mediate the effect of age on fitness

Despite older and female swallows with longer tail streamers exhibiting higher fecundity, appearing to be less social, and less social females exhibiting greater fecundity, we did not find evidence that social interactions mediated the relationship between age or tail streamer length and annual reproduction. In males, social interactions were not associated with any measure of paternity, a prerequisite for potential mediation in our analysis. In wild baboons (*Papio cynocephalus* and *P. anubis*) social bonds are not strong mediators of the relationship between exposure to adversity early in life and health and fitness outcomes [70,71]. Consistent in both species is a lack of mediation by social interactions, strong direct effects of the exposure to outcome, and notable independent effects of exposure to mediator and mediator to outcome. Unlike our results, positive selection on social connectivity has been observed in forked fungus beetles [13,72], although the relationship between social connectedness and fitness may vary depending on the size, age structure, and sex ratio of the group [73,74].

## Conclusion

Older barn swallows produced more offspring but whether this association reflected individual condition or selective removal requires further investigation. Our results do not support our original prediction that more social interactions with members of the opposite sex are part of the mechanistic pathway by which older individuals optimize mating opportunities to increase fitness. Rather, older birds seemed less social, and among females, less social birds had higher fecundity, possibly because they invested time in accruing resources instead of prospecting for mates. Our mediation models do not provide support for the hypothesis that social interactions are on the causal pathway between age and reproduction. A better understanding of how age, condition, and social interactions influence fitness requires future studies to track animals across their life course to assess within vs between individual change in these variables. Finally, it would be worth investigating whether social preference, assessed at the dyad instead of the node level, is a mechanism through which age and plumage traits affect reproductive success.

## Data and Code accessibility

All data and analysis code used in this paper are available at this this GitHub repository (https://github.com/laubach/bs_trait_dtrmnts_repro_success_mediation) and will be permanently archived at Zenodo upon final review.

## Authors’ contributions

Z.M.L., K.P.K., R.J.S. and I.L.L. conceptualized this study; I.L.L. R.J.S. and T.T. collected data; Z.M.L. conducted analyses with input from K.P.K.; Z.M.L. drafted the original manuscript; All authors reviewed and edited the drafts and gave final approval for publication.

## Acknowledgements

We thank Matt Aberle and Amanda Hund for help with fieldwork and Heather Kenny-Duddela for thoughtful feedback on early drafts of this manuscript. We thank Colorado Horse Rescue who gave us access to their property where barn swallows nest.

