## Supplemental Material for "Testing the mediating role of social interactions on the relationship between age and plumage traits on reproductive success"

**RESULTS**

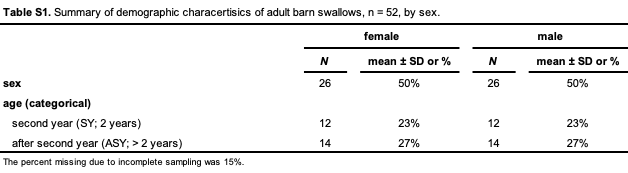

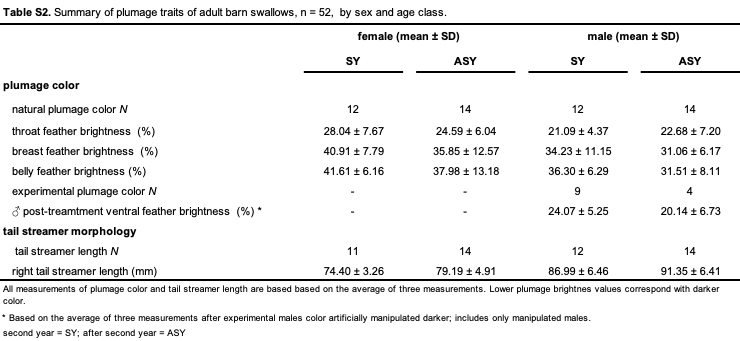

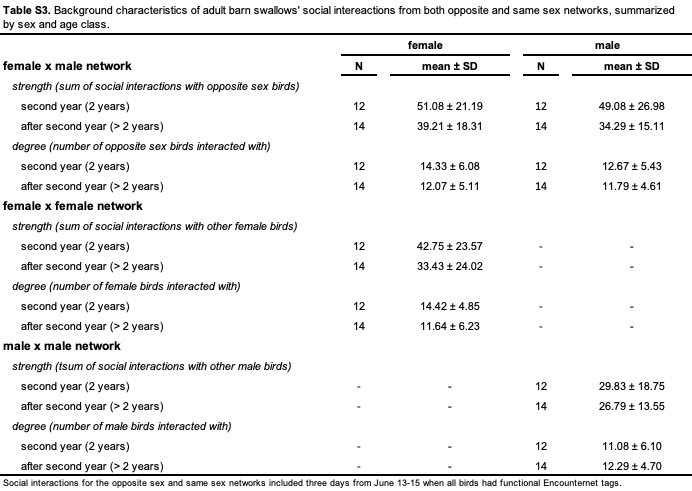

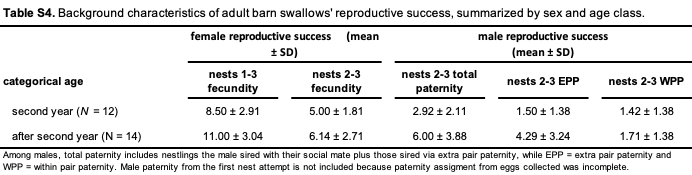

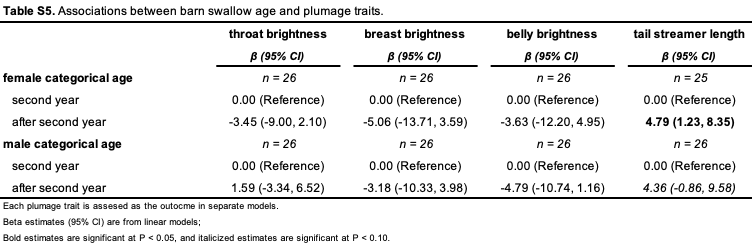

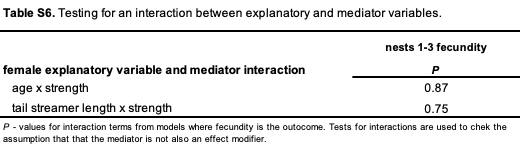

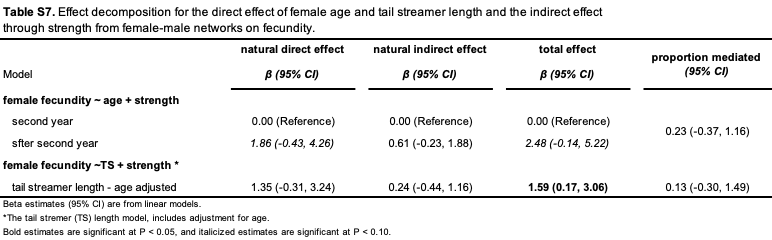
